# CCR5/CCL5-Dependent Mitochondrial Dysfunction Contributes to Angiotensin II– Induced Vascular Impairment in Mice

**DOI:** 10.64898/2026.03.10.710946

**Authors:** Gustavo Felix Pimenta, Ariane Bruder, Tyler Beling, Sergio Guerrero, Luis Oliveira de Moraes, Carlos Renato Tirapelli, Thiago Bruder-Nascimento

## Abstract

**AIM:** Chemokine signaling contributes to vascular inflammation and dysfunction in hypertension, yet the intracellular mechanisms linking CCL5/CCR5 activation to vascular impairment remain unclear. We tested the hypothesis that angiotensin II (Ang II) amplifies CCL5/CCR5 signaling to promote mitochondrial dysfunction and oxidative stress in the vasculature.

**METHODS:** Wild-type and CCR5-deficient mice were infused with Ang II for 14 days, and separate cohorts received recombinant CCL5. Vascular function and remodeling were assessed in aorta and mesenteric arteries, while mitochondrial respiration, membrane potential, and reactive oxygen species (ROS) production were evaluated in vascular smooth muscle cells (VSMCs).

**RESULTS:** Ang II increased circulating CCL5 levels and upregulated vascular CCR5 expression. CCR5 deficiency protected against Ang II–induced vascular dysfunction, remodeling, and inflammation. CCL5 infusion impaired endothelium-dependent relaxation and enhanced contractility without inducing structural remodeling. In VSMCs, CCL5 disrupted mitochondrial respiration, reduced maximal respiratory capacity, altered membrane potential, and increased mitochondrial ROS in a CCR5-dependent manner. Mitochondrial antioxidant treatment restored endothelial function but did not normalize enhanced contractility. In addition, vessels from CCL5-treated mice were unresponsive to acute mitochondrial uncoupling, consistent with impaired mitochondrial bioenergetic reserve.

**CONCLUSION:** Ang II amplifies CCL5/CCR5 signaling to drive mitochondrial dysfunction and oxidative stress, thereby promoting vascular impairment, and identify this pathway as a potential therapeutic target in hypertension.

## Introduction

Hypertension is a leading risk factor for cardiovascular morbidity and mortality and is characterized by vascular inflammation, endothelial dysfunction, and structural remodeling^1^. Angiotensin II (Ang II), a principal effector of the renin–angiotensin system, plays a central role in vascular injury by promoting oxidative stress, inflammatory signaling, vascular smooth muscle cell (VSMC) activation, and impaired endothelial nitric oxide bioavailability^2–4^. Although the hemodynamic effects of Ang II are well established, the inflammatory mediators that amplify and sustain Ang II–induced vascular dysfunction remain incompletely defined.

Chemokines and their receptors have emerged as critical regulators of vascular inflammation in hypertension. Among these, C-C motif ligand 5 (CCL5), also known as RANTES, and its receptor CCR5 have been implicated in immune cell recruitment and vascular injury^5–7^. Mikolajczyk et al. demonstrated that Ang II infusion increases CCL5 expression in perivascular adipose tissue and promotes the recruitment of CCR1-, CCR3-, and CCR5-expressing T lymphocytes to the vascular wall. Importantly, genetic deletion of CCL5 attenuated Ang II–induced perivascular inflammation, reduced vascular oxidative stress, and improved endothelial function, establishing a causal role for the CCL5 axis in hypertension-associated vascular dysfunction^8,9^. These findings identify CCL5 as a key mediator linking Ang II to vascular inflammation.

Beyond its role in immune cell recruitment, accumulating evidence suggests that CCR5 signaling exerts direct effects within the vascular wall^10^. We have previously shown that activation of the CCL5/CCR5 pathway promotes vascular remodeling and inflammatory gene expression^11^. Moreover, we demonstrated that aldosterone induces vascular dysfunction and hypertension through activation of the CCR5/CCL5 axis, identifying CCR5 as a functional mediator of mineralocorticoid-driven vascular injury^12^. Together, these findings support the concept that CCR5 is not merely a chemotactic receptor but an active participant in vascular pathophysiology.

Despite these advances, two major gaps remain. First, whether Ang II directly enhances CCL5/CCR5 signaling within vascular cells, thereby amplifying vascular responsiveness to inflammatory stimuli, is unclear. Second, the intracellular mechanisms by which CCL5/CCR5 activation promotes vascular dysfunction are not fully understood.

Mitochondrial dysfunction is increasingly recognized as a central driver of vascular disease^13–15^. Alterations in mitochondrial respiration, membrane potential, and redox balance contribute to endothelial dysfunction, abnormal VSMC contractility, and inflammatory signaling^16–18^. In addition to serving as a source of reactive oxygen species (ROS)^18^, mitochondrial bioenergetic status and membrane polarization regulate vascular tone by modulating ATP availability, calcium handling, and redox-sensitive pathways. Loss of respiratory reserve capacity and disruption of mitochondrial membrane potential can profoundly alter vascular reactivity and endothelial responses^19^. However, whether chemokine signaling through CCR5 directly modulates mitochondrial bioenergetics and membrane potential in VSMCs, thereby linking inflammatory signaling to vascular dysfunction, has not been investigated.

Based on our previous findings and the established role of CCL5 in Ang II–induced vascular inflammation, we hypothesized that Ang II amplifies CCL5/CCR5 signaling in the vasculature, leading to mitochondrial dysfunction characterized by altered respiration, disrupted membrane potential, and enhanced mitochondrial oxidative stress in VSMCs. We further hypothesized that these mitochondrial alterations contribute to endothelial dysfunction and abnormal vascular contractility. To test these hypotheses, we evaluated the impact of Ang II on the CCL5/CCR5 axis in vivo, assessed vascular injury in CCR5-deficient mice, and examined the direct effects of CCL5 on mitochondrial bioenergetics, redox balance, and vascular reactivity. Our findings identify a CCR5-dependent mitochondrial mechanism linking inflammatory chemokine signaling to vascular dysfunction.

## Methods

### Animals

Male wild-type (CCR5+/+, C57BL6/J) and global CCR5 knockout (CCR5−/−; Ccr5tm1Kuz/J) mice aged 10 and 14 weeks were used in this study. All animals were maintained on standard chow with ad libitum access to tap water. All mice were housed in an AAALAC-accredited facility within the College of Medicine at the University of South Alabama. Euthanasia was performed using CO₂ overdose. All procedures were approved by the Institutional Animal Care and Use Committee (IACUC) at the University of South Alabama (protocol #2219557) and conducted in accordance with the Guide for the Care and Use of Laboratory Animals.

### Angiotensin II Administration

Both CCR5+/+ and CCR5-/- mice were subjected to a 14-day infusion protocol using ALZET osmotic minipumps (Alzet Model 1002; Alzet Corp Durect, Cupertino, CA) to delivery ang II at a dose of 490 ng/min/kg^19^.

### CCL5 Administration

C57BL6/J male mice were subjected to a 14-day infusion protocol using ALZET osmotic minipumps (Alzet Model 1002; Alzet Corp Durect, Cupertino, CA) to delivery recombinant CCL5 at a dose of 0,42ng/day (CCL5 group) or saline (Control group).

### Plasma Analyses

Plasma levels of circulating chemokines and cytokines were analyzed using pooled samples from six mice per group as described before^12,20^.The analysis was performed with the Proteome Profiler Mouse Cytokine Array Kit (R&D Systems), following the manufacturer’s protocol. Band intensities were quantified using the ChemiDoc™ Imaging System (Bio-Rad Laboratories, Inc.), and the results were presented as fold changes in a heat map graph.

### Measurement of Plasma CCL5

Circulating CCL5 concentrations were determined in serum samples from CCR5+/+ mice treated with vehicle, Ang II and CCL5 using an enzyme-linked immunosorbent assay (ELISA) (R&D Systems).

### Real-time Polymerase Chain Reaction (RT-PCR)

mRNA was isolated from aortas, mesenteric arteries, perivascular adipose tissue (PVAT), and VSMCs using the RNeasy Mini Kit (Qiagen). Complementary DNA (cDNA) was synthesized by reverse transcription using SuperScript III (Thermo Fisher). Reverse transcription reactions were carried out at 58 °C for 50 minutes, followed by heat inactivation of the enzyme at 85 °C for 5 minutes. Gene expression was quantified by real-time quantitative RT-PCR using PowerTrack™ SYBR Green Master Mix (Thermo Fisher). Analyses were performed on a QuantStudio™ 5 Real-Time PCR System using 384-well plates (Thermo Fisher). Amplification was performed using gene-specific primers, as described in Supplementary Table 1 and 2. Relative gene expression was calculated using the 2ΔΔ Ct method and expressed as fold change, indicating upregulation or downregulation of gene expression.

### Vascular Reactivity Studies

Segments of thoracic aorta and mesenteric arteries from CCR5+/+ and CCR5−/− mice treated with vehicle or Ang II, as well as from CCR5+/+ mice treated with vehicle or CCL5, were isolated, cut into 2-mm rings, and mounted in a wire myograph for isometric tension recordings. Experiments were performed in Krebs–Henseleit solution maintained at 37 °C and continuously aerated with 95% O₂ and 5% CO₂, with the following composition (in mM): 130 NaCl, 4.7 KCl, 1.17 MgSO₄, 0.03 EDTA, 1.6 CaCl₂, 14.9 NaHCO₃, 1.18 KH₂PO₄, and 5.5 glucose. After 30 minutes of stabilization, vascular viability was assessed by KCl-induced contraction (60 mM). Concentration–response curves were then generated for phenylephrine (PE), acetylcholine (ACh), and sodium nitroprusside (SNP). Some experiments were performed in the presence of MnTMPyP (30 µM), a mitochondrial antioxidant and superoxide dismutase 2 (SOD2) mimetic, or carbonyl cyanide *m*-chlorophenyl hydrazone (CCCP, 1 µM), a mitochondrial uncoupler. Maximal response (Emax) and the negative logarithm of the half-maximal effective concentration (pD2) were calculated were analyzed.

### Aortic Remodeling

Thoracic aortas were harvested from CCR5+/+ and CCR5−/− mice treated with vehicle or Ang II, and from CCR5+/+ mice treated with vehicle or CCL5. Following fixation in 4% paraformaldehyde for 12 h and storage in 70% ethanol, tissues were paraffin-embedded, serially sectioned, and stained with Masson’s trichrome to assess vascular structure. Morphometric analysis was performed to media-to-lumen ratio (M/L ratio), medial cross-sectional area (CSA), and medial thickness.

### Mesenteric Artery Remodeling

Mesenteric arteries were isolated from vehicle or CCL5-treated mice and mounted in a pressure myograph system. After equilibration at physiological pressure, internal and external diameters, wall thickness, lumen area, and media-to-lumen ratio were determined at 60 mmHg. Media-to-lumen ratio was further evaluated over a pressure range of 10–160 mmHg, and vascular mechanical properties were assessed by calculating stress–strain relationships.

### Perivascular Adipose Tissue (PVAT) Incubation

Mesenteric PVAT from wild-type mice was isolated and incubated in vitro for 24 hours with Ang II (0.1 μM) at 37°C. Vehicle-treated tissues were used as controls. After incubation, total RNA was extracted, and gene expression was assessed by real-time quantitative PCR (RT-PCR).

### Vascular Smooth Muscle Cell (VSMCs) Experiments

Rat aortic smooth muscle cells (RASMCs; Lonza, Walkersville, MD, USA) were cultured in Dulbecco’s Modified Eagle Medium (DMEM; Gibco, Thermo Fisher Scientific, Waltham, MA, USA) supplemented with 10% fetal bovine serum (FBS; HyClone, Logan, UT, USA), 100 U/mL penicillin, and 100 µg/mL streptomycin, as previously described^11^. Cells were maintained at 37°C in a humidified atmosphere containing 5% CO₂, and the culture medium was replaced every 2–3 days. Prior to stimulation, cells were serum-starved (0.5-1% FBS) for 18-24 h to minimize basal signaling and synchronize cellular responses.

RASMCs were treated with vehicle, CCL5, or Ang II (0.1 μM), either alone or in combination with specific pharmacological blockers, according to the experimental protocol. When applicable, inhibitors were added 30 minutes before agonist stimulation.

### Oxygen Consumption Rate (OCR) Analysis

VSMCs were treated with vehicle or CCL5, and mitochondrial respiration was assessed using a Seahorse XF extracellular flux analyzer (Agilent Technologies) with XF24 microplates as described before^19^. Cells were seeded onto Seahorse plates and incubated overnight. On the following day, the culture medium was replaced with Seahorse assay medium, and cells were equilibrated in a non-CO₂ incubator at 37 °C. Oxygen consumption rate (OCR) was determined under basal conditions and after sequential injections of oligomycin, FCCP, and rotenone/antimycin To specifically evaluate mitochondrial respiration, OCR values were corrected for non-mitochondrial respiration using the rotenone/antimycin A measurements. Data were normalized to protein content^21^.

### Assessment of Mitochondrial Membrane Potential (JC-1 Assay)

VSMCs were treated with CCL5 and incubated with the mitochondrial membrane potential-sensitive dye JC-1 (Invitrogen). Cells were incubated with JC-1 diluted in prewarmed HBSS at a final concentration of 2 µM for 30 minutes at 37 °C in the dark. After incubation, cells were washed with PBS and analyzed by fluorescence microscopy. Mitochondrial membrane potential was assessed by the ratio of red (J-aggregates) to green (J-monomers) fluorescence^19^.

### MitoSOX Fluorescence Analysis

VSMCs were treated with CCL5 and subsequently incubated with MitoSOX™ Red (Invitrogen, Thermo Fisher Scientific, Waltham, MA, USA) to assess mitochondrial superoxide production. Cells were incubated with MitoSOX diluted in prewarmed HBSS at a final concentration of 100 nM for 10 minutes at 37 °C in the dark. After incubation, cells were washed with PBS and analyzed by fluorescence microscopy^19^.

### Hydrogen Peroxide Measurement (Amplex Red Assay)

VSMCs were treated with CCL5 in the presence or absence of MnTMPyP (30 µM)^22^ and subsequently collected in lysis buffer consisting of Hank’s Balanced Salt Solution supplemented with protease inhibitors (Complete Mini) and phosphatase inhibitors (PhosSTOP). Cell lysis was performed by five consecutive freeze–thaw cycles, followed by passage of the lysate through a 30-gauge needle five times to ensure complete membrane disruption for subsequent determination of reactive oxygen species (ROS), as previously described^12^. The homogenates were then centrifuged at 1,000 × g for 5 minutes at 4 °C to remove intact cells, nuclei, and cellular debris. Throughout the entire procedure, samples were maintained at temperatures close to 0 °C. RASMC lysates were subsequently incubated in Amplex Red assay buffer containing 25 mM HEPES (pH 7.4), 120 mM NaCl, 3 mM KCl, and 1 mM MgCl₂, supplemented with 0.1 mM Amplex Red (Thermo Scientific, #A22188, Massachusetts, USA) and 0.35 U/mL horseradish peroxidase (HRP), in the presence or absence of catalase (300 U/mL). The reaction was initiated by the addition of NADPH (36 µM). Fluorescence was measured using a Synergy 4 multimode microplate reader (Biotek), with excitation at 530/25 nm and emission at 590/35 nm, at 25 °C for 1 hour.

### MitoTracker Red Staining

VSMCs were treated with CCL5 and subsequently incubated with MitoTracker probes (Invitrogen, Thermo Fisher Scientific, Waltham, MA, USA) to assess mitochondrial mass and activity. Cells were incubated with MitoTracker Green and/or MitoTracker Red, diluted in prewarmed HBSS culture medium, at a final concentration of 100 nM for 30 minutes at 37 °C in the dark. After incubation, cells were washed with PBS and analyzed by fluorescence microscopy.

### Western blotting

Proteins from vascular smooth muscle cells were extracted and directly homogenized in 2× Laemmli sample buffer supplemented with 2-mercaptoethanol (β-mercaptoethanol) (Bio-Rad, Hercules, CA, USA). Protein samples were separated by SDS– polyacrylamide gel electrophoresis using gradient gels (Bio-Rad) and subsequently transferred onto poly (vinylidene difluoride) (PVDF) membranes (Immobilon-P). Membranes were blocked with either 5% non-fat dry milk or 1% bovine serum albumin (BSA) prepared in Tris-buffered saline containing Tween-20 (TBS-T) for 1 h at room temperature. Membranes were then incubated overnight at 4 °C with specific primary antibodies, as described in Supplementary Table 3. After incubation with the appropriate secondary antibodies, protein bands were detected using an enhanced luminol-based chemiluminescence reagent (SuperSignal™ West Femto Maximum Sensitivity Substrate, Thermo Scientific, No. 34095, Massachusetts, USA).

### Statistical Analysis

Comparisons among four experimental groups were performed using two-way analysis of variance (ANOVA), followed by Tukey’s multiple-comparison test. Differences between two groups were assessed using Student’s *t*-test. Concentration-response curves were fitted by nonlinear regression to evaluate the relationship between agonist concentrations and the resulting contractile or relaxant responses. All statistical analyses were carried out using GraphPad Prism version 8.0 (GraphPad Software Inc., San Diego, CA).

## Results

### Ang II enhances CCL5/CCR5 signaling

We first examined whether Ang II treatment amplifies components of the CCL5/CCR5 signaling axis in the circulation and vasculature. Proteomic profiling of plasma from Ang II–treated mice revealed a pronounced imbalance in cytokine and chemokine levels, with a notable increase in CCL5 (Fig. 1A). Elevated circulating CCL5 levels were subsequently confirmed by ELISA (Fig. 1B). We next assessed the expression of CCL5 and its receptors, CCR1, CCR3, and CCR5, and CCR2 in vascular tissues. In the aorta, Ang II significantly increased the gene expression of CCL5 as well as CCR2, CCR3, and CCR5 (Fig. 1C). In contrast, in mesenteric arteries, Ang II increased CCL5 and CCR5 expression (Fig. 1D). Finally, to determine whether Ang II directly modulates this pathway in VSMCs, cells were treated with Ang II and the expression of CCL5, and its receptors was assessed. Ang II significantly increased the expression of CCR5, as well as CCR1 and CCR3, while CCL5 expression remained unchanged (Fig. 1E). These findings indicate that Ang II directly enhances the responsiveness of VSMCs to CCL5 by upregulating its receptors, thereby amplifying CCL5/CCR5 signaling.

**Figure 1.**
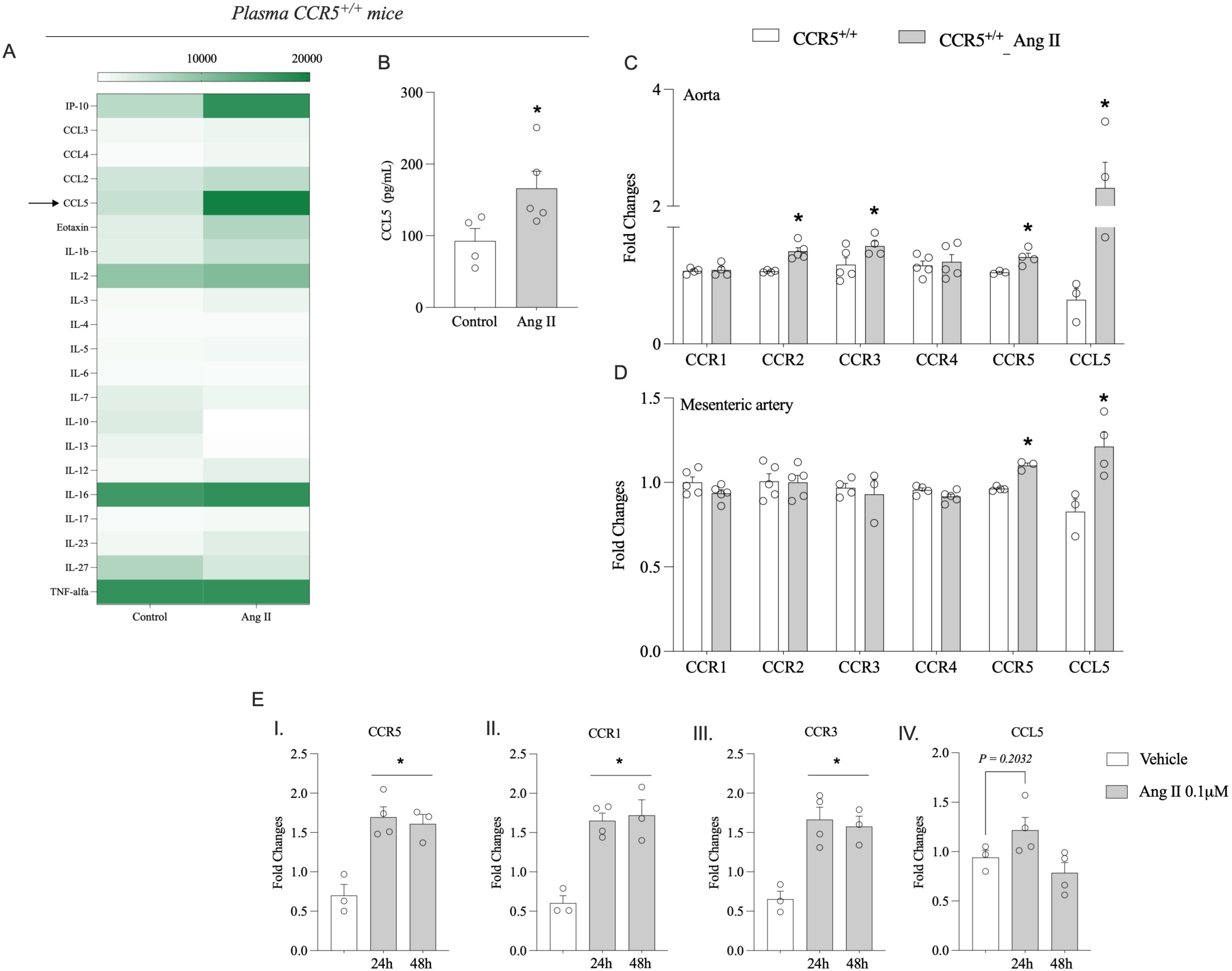
Ang II enhances CCL5/CCR5 signaling in the circulation and vasculature. Inflammatory profile in plasma from wild-type (WT) mice infused with vehicle (saline) or angiotensin II (Ang II; 490 ng/kg/min for 14 days via osmotic minipump), assessed using a Proteome Profiler Mouse Cytokine Array and presented as a heat map (A; pooled samples from 6 mice per group). Circulating CCL5 levels measured by ELISA (B). Gene expression of CCL5 and its receptors in thoracic aortae (C) and mesenteric arteries (D) from WT mice infused with vehicle or Ang II, determined by real-time RT-PCR. Chemokine receptor expression in rat aortic smooth muscle cells (RASMCs) treated with vehicle or Ang II (0.1 µM) for 24 h was assessed by real-time RT-PCR (E). Data are presented as mean ± SEM, (n= 3-5). Statistical analysis was performed using Student t test and one-way ANOVA followed by Tukey’s post hoc test. *P < 0.05 vs. vehicle.

### Ang II induces vascular injury via CCR5

We first characterized the effects of Ang II on vascular function in the aorta and mesenteric arteries, as well as on aortic remodeling and vascular inflammation. Ang II treatment induced marked hypercontractility to phenylephrine in both aortic (Fig. 2A) and mesenteric rings (Fig. 2B). Given the important role of PVAT in modulating vascular tone, we next assessed mesenteric arteries with intact PVAT and found that Ang II abolished the anti-contractile effects of PVAT (Supplementary Fig. 1A-B). In addition, Ang II promoted aortic remodeling (Fig. 2C), characterized by ratio (M/L ratio) (I), medial cross-sectional area (CSA) (II) and medial thickness (III). Ang II also increased vascular inflammation in the aorta, as evidenced by elevated ICAM-1, CCL2, and VCAM-1 expression (Fig. 2D), and in mesenteric arteries, as indicated by increased ICAM-1, F4/80, IL-6, CCL2, and IL-16 (Fig. 2E).

**Figure 2.**
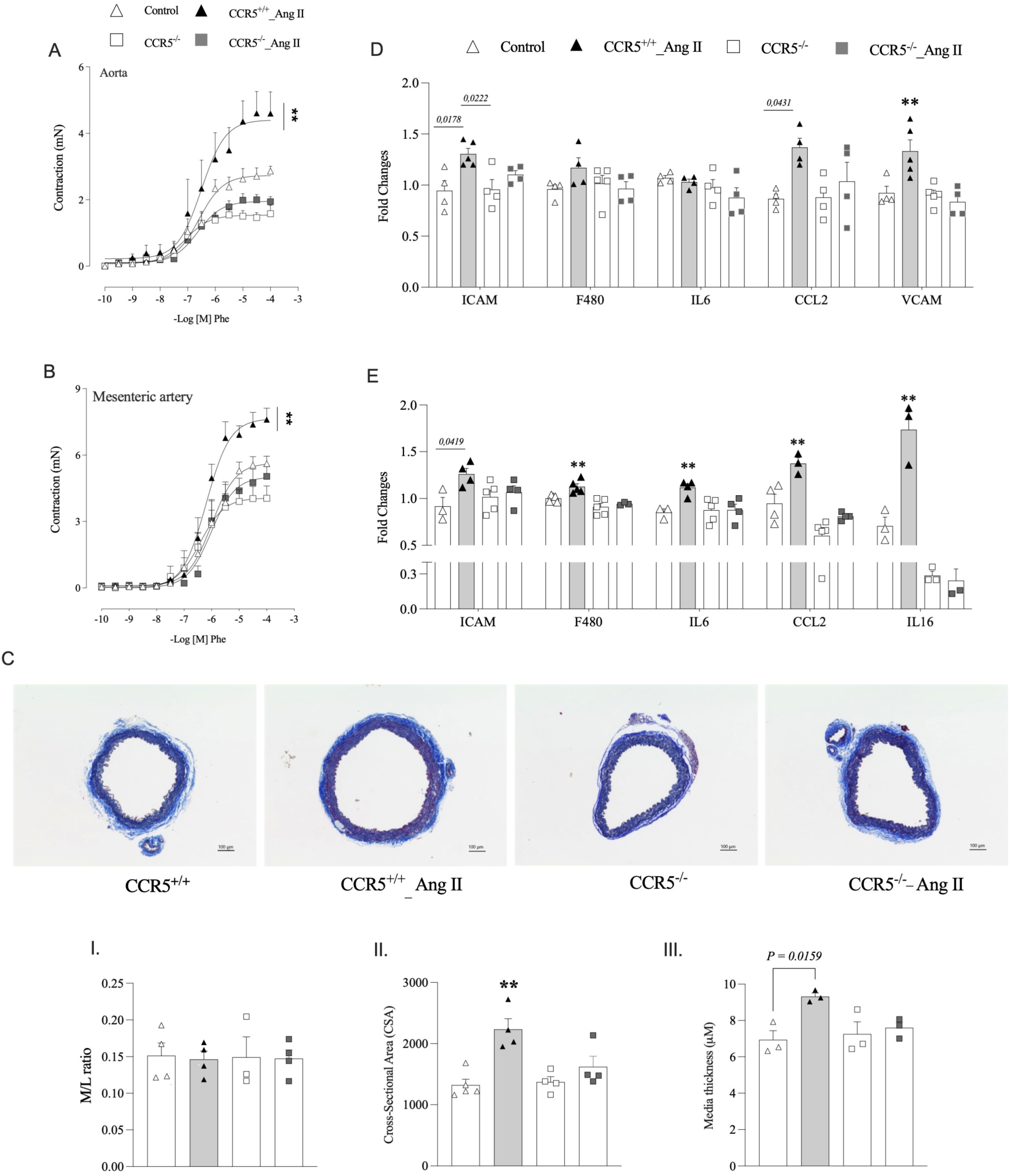
CCR5 deficiency protects against Ang II–induced vascular dysfunction and remodeling. Concentration–response curves to phenylephrine (Phe) in thoracic aortae (A) and mesenteric arteries (B) from wild-type (WT) and CCR5⁻/⁻ mice treated with vehicle or angiotensin II (Ang II; 490 ng/kg/min for 14 days via osmotic minipump). Representative Masson’s trichrome–stained sections of thoracic aorta are shown (C, top), with corresponding quantification of ratio (M/L ratio) (I), medial cross-sectional area (CSA) (II) and medial thickness (III) presented as bar graphs (bottom). Gene expression analysis of inflammatory markers in aortae (D) and mesenteric arteries (E) was performed by real-time RT-PCR. Data are expressed as mean ± SEM (n = 3–5 per group). Emax and pD₂ values were calculated by nonlinear regression. Statistical analysis was performed using two-way ANOVA followed by Tukey’s post hoc test. **P < 0.05 vs. Control; CCR5⁻/⁻ and CCR5⁻/⁻_ Ang II (Emax).

To determine the contribution of CCR5 to these effects, we used CCR5-deficient (CCR5⁻/⁻) mice. Genetic deletion of CCR5 conferred protection against Ang II–induced vascular dysfunction, remodeling, and inflammation (Fig. 2A-E). Similarly, Ang II– associated PVAT dysfunction was also reversed in CCR5⁻/⁻ mice, indicating that CCR5 contributes to both vascular and PVAT alterations induced by Ang II. However, PVAT from mesenteric arteries incubated with Ang II displayed increased CCR3 and VCAM-1, with no difference in CCR5 and CCL5 (Supplementary Fig. 1C), therefore, for the following studies, we only focused on vessels without PVAT.

### CCL5 induces vascular dysfunction and inflammation

We previously demonstrated that CCL5 promotes vascular dysfunction in vitro and ex vivo^11,23^; here, we extend these findings to an in vivo setting. WT mice were infused with recombinant CCL5 at a dose selected to replicate the circulating CCL5 levels observed following Ang II treatment. At this dose (0,42ng/day), circulating CCL5 concentrations closely matched those measured in Ang II–treated mice (Fig. 3A) and were therefore used for subsequent experiments.

**Figure 3.**
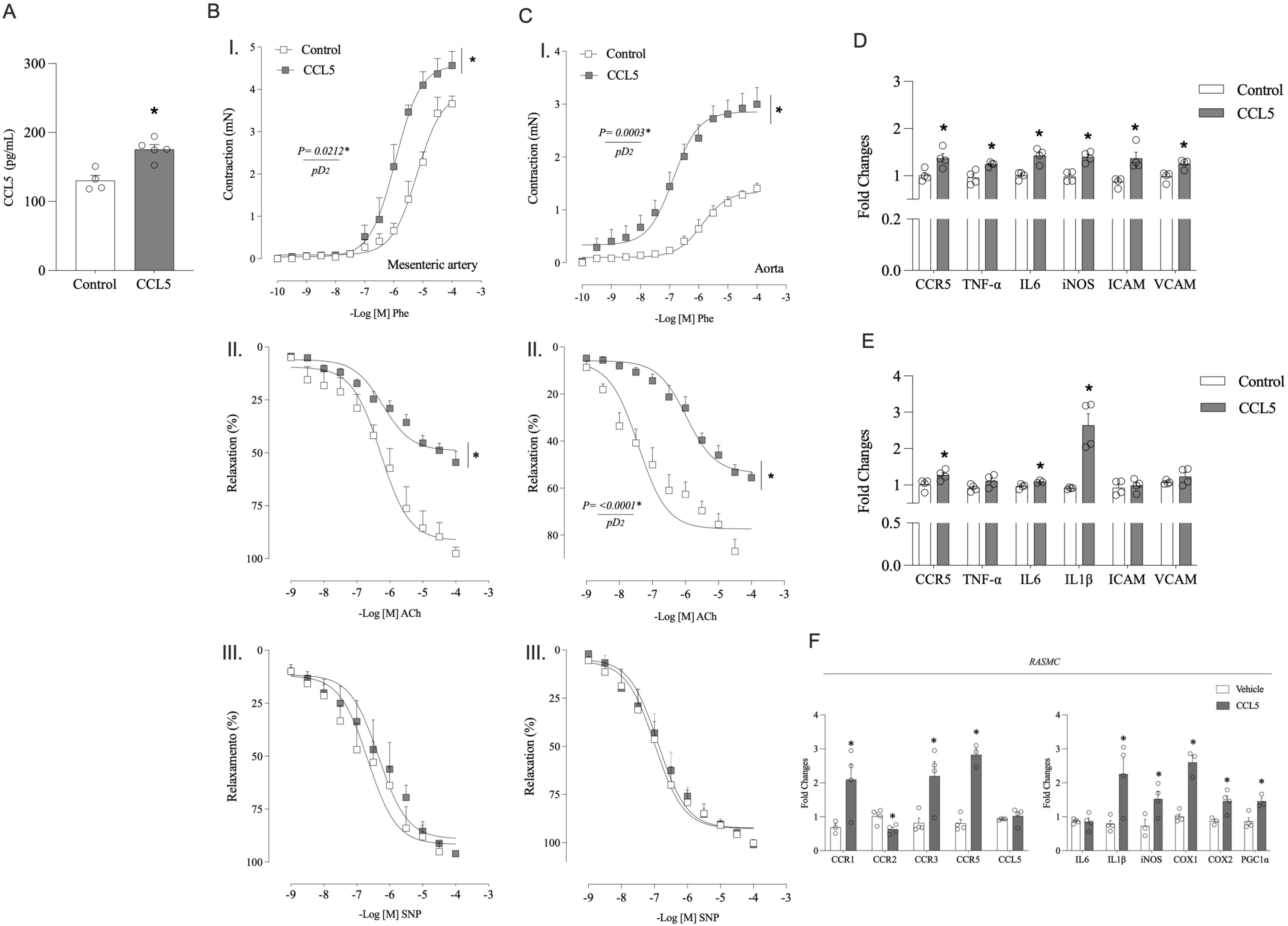
CCL5 induces vascular dysfunction and inflammation. Plasma CCL5 levels measured by ELISA in wild-type mice infused with vehicle (saline) or recombinant CCL5 (0.42 ng/day for 14 days via osmotic minipump) (A). Concentration–response curves in mesenteric arteries (B) to phenylephrine (Phe; I), acetylcholine (ACh; II), and sodium nitroprusside (SNP; III). Concentration–response curves in thoracic aortae (C) to Phe (I), ACh (II), and SNP (III). Gene expression of inflammatory markers in mesenteric arteries (D) and aortae (E), determined by real-time RT-PCR. Gene expression in rat aortic smooth muscle cells (RASMCs) treated with vehicle or CCL5 was assessed by real-time RT-PCR (F). Data are expressed as mean ± SEM (n = 3–5 per group). Emax and pD₂ values were calculated by nonlinear regression. Statistical comparisons were performed using Student’s t test. *P < 0.05 vs. Vehicle (Emax).

In mesenteric arteries (Fig. 3B), CCL5 treatment significantly altered vascular reactivity, characterized by enhanced contractile responses to Phe (I) and impaired endothelium-dependent relaxation to Ach (II), while endothelium-independent relaxation to SNP remained unaffected (III). Similarly, in the aorta (Fig. 3C), CCL5 increased vascular contractility (I) and reduced endothelium-dependent relaxation to ACh (II) without altering responses to SNP (III). These findings demonstrate that CCL5 induces vascular dysfunction in both resistance and conduit vessels.

Consistent with these functional impairments, gene expression analyses revealed increased expression of inflammatory markers in both vascular beds following CCL5 treatment, indicating an enhanced vascular inflammatory response (Fig. 3D-E). In contrast, CCL5 treatment did not induce aortic or mesenteric remodeling (Supplementary Fig. 2), as assessed by histological analysis (Fig. 2A). and pressure myography (Fig. 2B).

Finally, to determine whether CCL5 directly affects VSMCs, RASMC were treated with CCL5. CCL5 significantly upregulated CCL5 receptors (CCR1, CCR3 and CCR5) and increased the expression of inflammatory genes in VSMCs, supporting a direct pro-inflammatory effect of CCL5 on VSMCs (Fig. 3F).

### CCL5-Induced Vascular Dysfunction Is Mediated by Mitochondrial Dysfunction

Mitochondrial dysfunction is a key contributor to vascular injury. To determine whether CCL5 induces mitochondrial dysfunction in VSMCs, mitochondrial respiration was assessed by measuring the oxygen consumption rate (OCR) in RASMC treated with CCL5 for 24 hours. Representative OCR traces demonstrated that CCL5 altered cellular respiration compared with vehicle-treated cells (Fig. 4A). To specifically evaluate mitochondrial function, OCR values were corrected by subtracting rotenone-plus antimycin A–insensitive respiration (Fig. 4B). Quantitative analysis revealed a significant increase in basal mitochondrial respiration (I), no change in ATP-linked respiration (II), a marked reduction in maximal respiratory capacity (III) and reserve capacity (IV), indicating increased basal respiratory demand and diminished respiratory reserve. These alterations were accompanied by an increase in MitoTracker Red fluorescence, indicating changes in mitochondrial membrane potential and/or mitochondrial network organization in response to CCL5 (Fig. 4C I–II). Representative fluorescence images demonstrate enhanced mitochondrial signal in CCL5-treated VSMCs compared with control cells, consistent with altered mitochondrial polarization and remodeling rather than a direct measure of mitochondrial mass.

**Figure 4.**
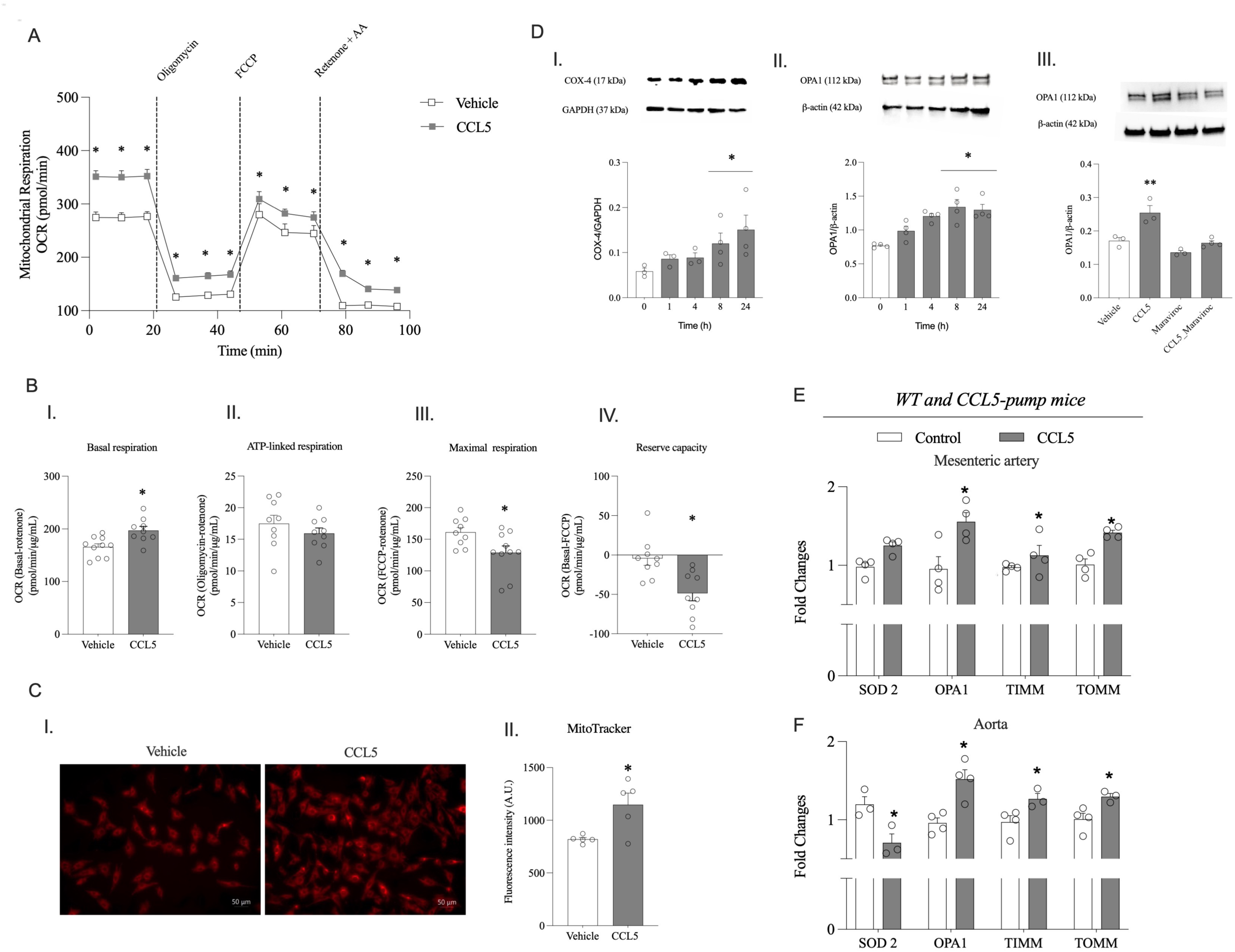
CCL5 impairs mitochondrial respiration and promotes mitochondrial remodeling in VSMCs. Representative oxygen consumption rate (OCR) trace over time in rat aortic smooth muscle cells (RASMCs) treated with vehicle or CCL5 for 24 h (A). Quantification of mitochondrial respiration parameters (B), including basal respiration (I), ATP-linked respiration (II), and maximal respiration (III). To specifically assess mitochondrial respiration, OCR values were corrected by subtracting rotenone- and antimycin A–insensitive (non-mitochondrial) respiration. Mitochondrial content and/or membrane polarization were evaluated using MitoTracker staining (C), showing representative fluorescence images (I) and quantification of MitoTracker fluorescence intensity (II). Western blot analysis of mitochondrial remodeling proteins in RASMCs treated with CCL5 (D): time-course expression of COX IV (I) and OPA1 (II), and the effect of CCR5 antagonism with maraviroc on CCL5-induced OPA1 expression (III). Gene expression of mitochondrial dynamics and redox-related genes in mesenteric arteries (E) and thoracic aortae (F) from mice infused with vehicle or CCL5 for 14 days, determined by real-time RT-PCR. Data are expressed as mean ± SEM (n = 3–5 per group). Statistical analyses were performed using Student’s t test or two-way ANOVA followed by Tukey’s post hoc test, as appropriate. *P < 0.05 vs. Vehicle; **P < 0.05 vs. Vehicle and CCL5 + Maraviroc.

In agreement with these observations, CCL5 induced changes in proteins associated with mitochondrial remodeling (Fig. 4D). Time-course analysis revealed increased expression of COX-4 (I) and the fusion-related protein OPA1 (II), beginning 8 hours after CCL5 exposure. Importantly, pharmacological antagonism of CCR5 with maraviroc prevented CCL5-induced upregulation of OPA1 (III), indicating that mitochondrial remodeling is mediated through CCR5 signaling.

Supporting the persistence of these mitochondrial alterations in vivo, 14 days of CCL5 treatment modified the expression of genes involved in mitochondrial dynamics, remodeling, and antioxidant defense in vascular tissues, including SOD2, OPA1, TIMM, and TOMM, in mesenteric arteries and aorta (Fig. 4E–F).

### CCL5 enhances mitochondrial oxidative stress and alters redox balance

Given the observed mitochondrial dysfunction, we next examined whether CCL5 promotes mitochondrial oxidative stress. CCL5 significantly increased mitochondrial superoxide production in VSMCs, as evidenced by enhanced MitoSOX fluorescence (Fig. 5A). Importantly, this effect was abolished by pharmacological blockage of CCR5 with maraviroc, indicating that CCL5-induced mitochondrial ROS generation is CCR5-dependent. In addition, CCL5 elevated intracellular H₂O₂ levels, an effect that was prevented by treatment with the mitochondrial-targeted antioxidant MnTMPyP (Fig. 5B).

**Figure 5.**
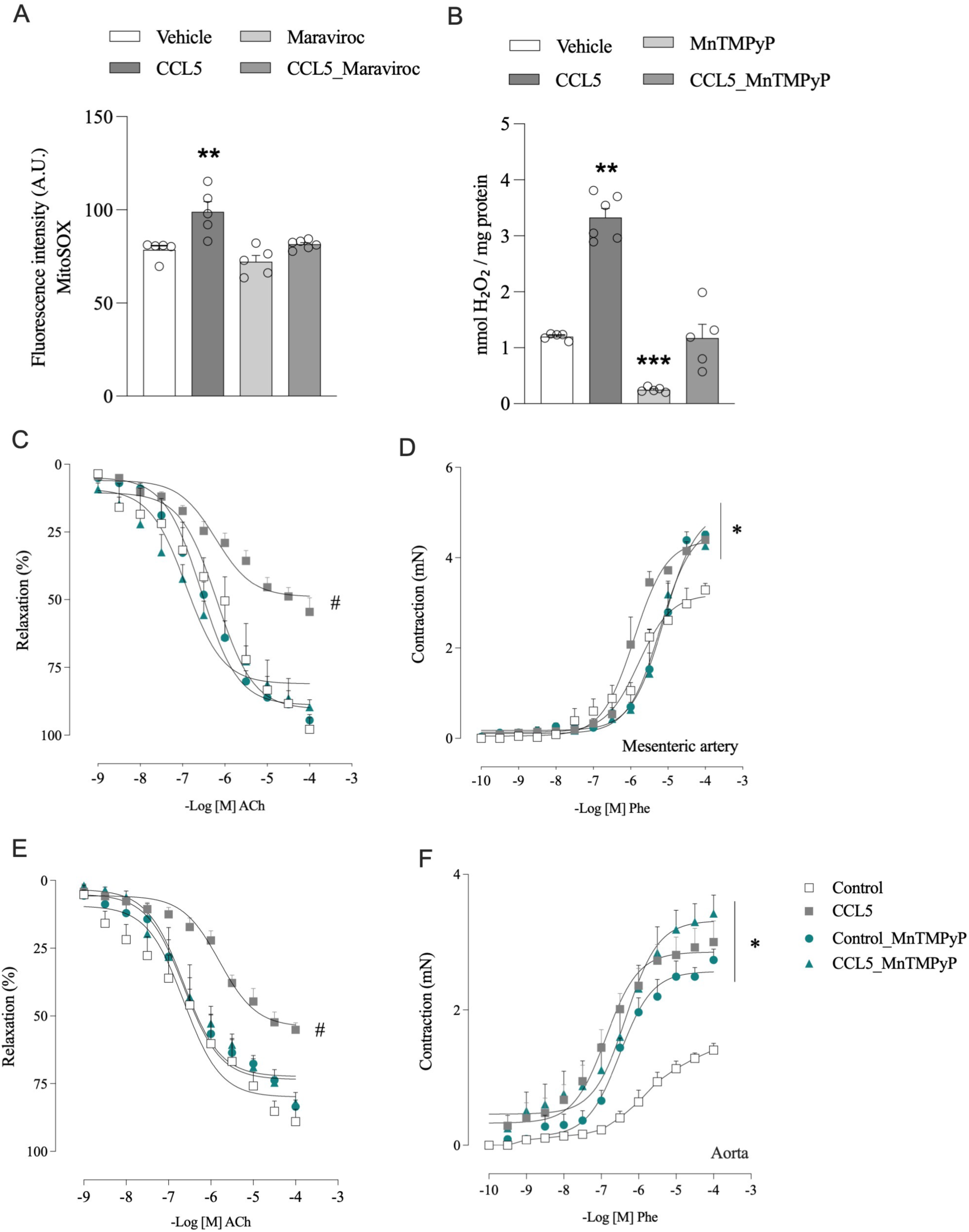
CCL5 enhances mitochondrial oxidative stress and mitochondrial ROS contributes to endothelial dysfunction. Mitochondrial superoxide production in rat aortic smooth muscle cells (RASMCs), assessed by MitoSOX fluorescence (A), and intracellular H₂O₂ levels measured by Amplex Red assay (B) following treatment with vehicle or CCL5, in the presence or absence of the CCR5 antagonist maraviroc. Concentration–response curves in mesenteric arteries to acetylcholine (ACh; C) and phenylephrine (Phe; D), and in thoracic aortae to ACh (E) and Phe (F), from mice treated with vehicle or CCL5, evaluated in the presence or absence of the mitochondrial-targeted antioxidant MnTMPyP. Data are expressed as mean ± SEM (n = 5–6 per group). Emax and pD2 values were calculated by nonlinear regression. Statistical analysis was performed using two-way ANOVA followed by Tukey’s post hoc test. *P < 0.05 vs. control (Emax); **P < 0.05 vs. vehicle, maraviroc, and CCL5_maraviroc. **P < 0.05 vs. vehicle, MnTMPyP, and CCL5_MnTMPyP; ***P < 0.05 vs. vehicle, CCL5, and CCL5_maraviroc. ^#^P < 0.05 vs. control, control_MnTMPyP, and CCL5_MnTMPyP (Emax).

To determine whether mitochondrial oxidative stress contributes to vascular dysfunction, vascular reactivity was assessed in the presence of the MnTMPyP. In mesenteric arteries from CCL5-treated animals, MnTMPyP restored endothelium-dependent relaxation to ACh (Fig. 5C) but did not normalize the augmented contractile responses to Phe (Fig. 5D). Similar results were observed in aortic rings, where MnTMPyP improved ACh-induced relaxation (Fig. 5E) without attenuating the enhanced Phe-induced contraction (Fig. 5F). Notably, MnTMPyP increased vasoconstrictor responses in vessels from control animals.

These findings indicate that mitochondrial ROS is a key mediator of CCL5-induced endothelial dysfunction. In contrast, the enhanced vascular contractility appears to be largely independent of mitochondrial ROS under these conditions.

### CCL5 disrupts mitochondrial membrane potential and promotes mitochondrial remodeling via CCR5

Given the impairment in mitochondrial respiration, we next evaluated mitochondrial membrane potential. JC-1 fluorescence analysis demonstrated mitochondrial depolarization in CCL5-treated VSMCs (Fig. 6A), evidenced by reduced aggregate (red) fluorescence (II), increased monomer (green) fluorescence (III), and a decreased red/green fluorescence ratio compared with vehicle-treated cells (IV).

**Figure 6.**
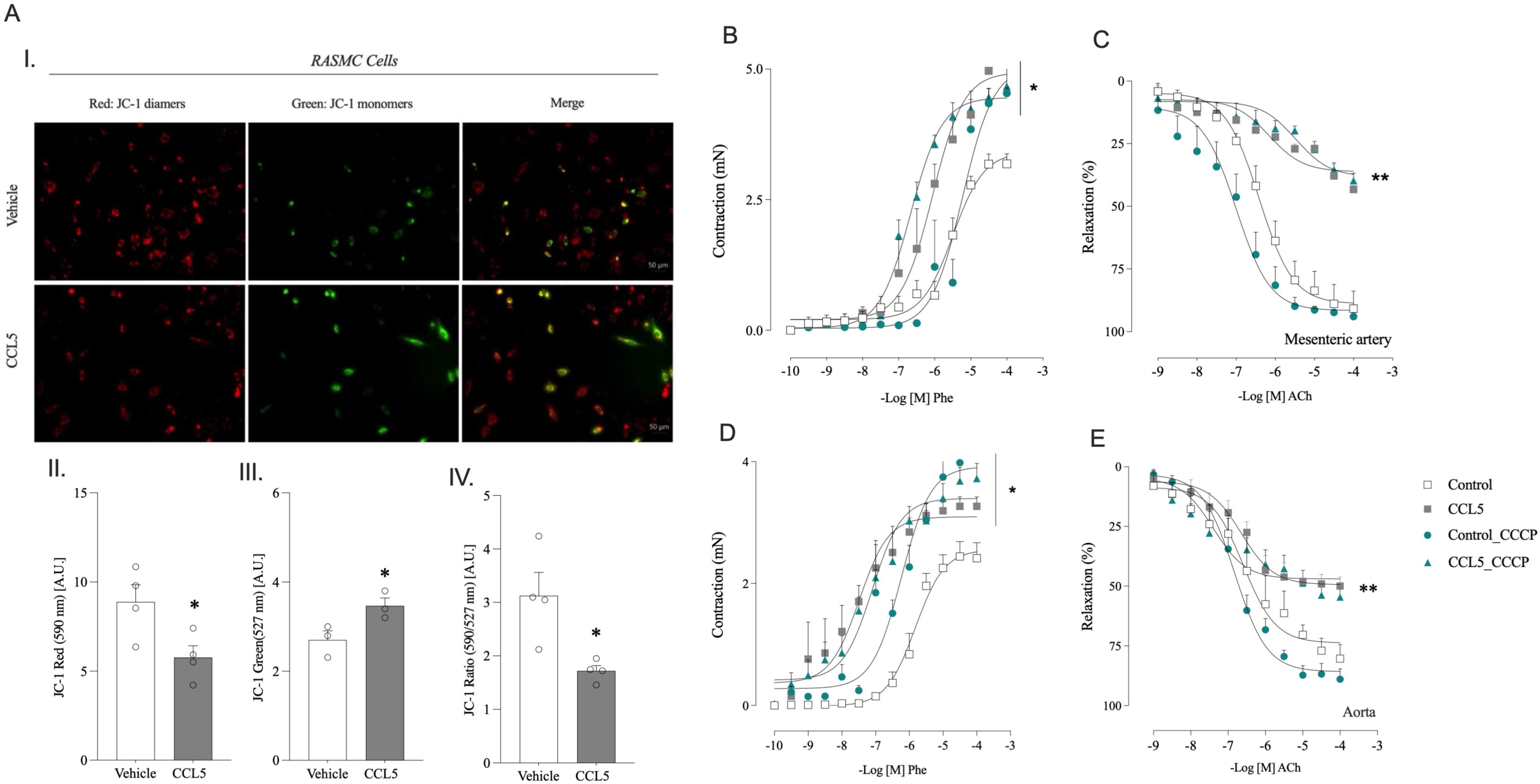
CCL5 disrupts mitochondrial membrane potential and alters vascular responses to mitochondrial uncoupling. Mitochondrial membrane potential (ΔΨm) in rat aortic smooth muscle cells (RASMCs) treated with vehicle or CCL5, assessed by JC-1 staining (A). Representative fluorescence images are shown (I), along with quantification of JC-1 aggregate (red) fluorescence (II), JC-1 monomer (green) fluorescence (III), and the red/green fluorescence ratio (IV), reflecting mitochondrial membrane potential. Concentration–response curves in mesenteric arteries to phenylephrine (Phe; B) and acetylcholine (ACh; C), and in thoracic aortae to Phe (D) and ACh (E), from mice treated with vehicle or CCL5, evaluated in the presence or absence of the mitochondrial uncoupler CCCP. Data are expressed as mean ± SEM (n = 3–5 per group). Emax and pD2 values were calculated by nonlinear regression. Statistical analysis was performed using two-way ANOVA followed by Tukey’s post hoc test, while other comparisons were performed using Student’s t-test, as appropriate. *P < 0.05 vs. vehicle or control; **P < 0.05 vs. control and control_CCCP (Emax).

Supporting these in vitro findings, Ang II treatment in vivo altered the expression of mitochondrial dynamics–related genes in both aorta and mesenteric arteries, effects that were prevented in CCR5-deficient mice (Supplementary Fig. 3A–B).

To further assess the contribution of mitochondrial bioenergetics to vascular reactivity, vessels were exposed to the mitochondrial uncoupler CCCP. In arteries from control animals (not treated with CCL5), CCCP impaired vascular responses, indicating that mitochondrial membrane potential contributes to the regulation of vascular tone under physiological conditions. In contrast, CCCP did not modify the vascular dysfunction observed in CCL5-treated preparations (Fig. 6B), including endothelial dysfunction (Fig. 6C). Similar results were obtained in aortic rings, where CCCP failed to attenuate Phe-induced contraction (Fig. 6D) or improve ACh-induced relaxation (Fig. 6E).

These findings suggest that chronic CCL5 exposure induces sustained disruption of mitochondrial membrane potential and bioenergetic reserve, rendering the vasculature insensitive to further acute mitochondrial uncoupling. Together, the data support distinct mitochondrial mechanisms underlying endothelial dysfunction and enhanced vascular contractility following CCL5 stimulation.

## Discussion

The present study identifies a CCR5-dependent mitochondrial mechanism linking Ang II–induced chemokine signaling to vascular dysfunction. We demonstrate that Ang II enhances CCL5/CCR5 signaling in the vasculature, that CCR5 deficiency protects against Ang II–induced vascular hypercontractility, remodeling, and inflammation, and that CCL5 directly disrupts mitochondrial respiration, reduces respiratory reserve capacity, alters membrane potential, and increases mitoROS in VSMCs. Importantly, mitochondrial ROS selectively mediated endothelial dysfunction, whereas enhanced vascular contractility was associated with sustained disruption of mitochondrial bioenergetics and membrane potential. These findings establish mitochondrial dysfunction as a central downstream effector of CCR5 signaling in vascular pathology.

Chemokine-driven vascular inflammation has been previously implicated in Ang II– induced hypertension. Mikolajczyk et al^9^. demonstrated that Ang II infusion increases CCL5 expression in PVAT and promotes recruitment of chemokine receptor–expressing T lymphocytes to the vessel wall; deletion of CCL5 attenuated vascular oxidative stress and preserved endothelial function. Our findings extend this paradigm by showing that Ang II not only increases circulating CCL5 but also upregulates CCR5 expression within the vascular wall and in VSMCs, thereby amplifying responsiveness to chemokine signaling. Moreover, CCR5 deficiency conferred protection against Ang II–induced vascular dysfunction and remodeling, highlighting CCR5 as a functional mediator of Ang II–driven vascular injury.

These results build upon our previous work demonstrating a pathogenic role for the CCL5/CCR5 axis in vascular disease^10–12,24^. We previously showed that CCL5/CCR5 activation promotes vascular remodeling and inflammatory signaling^11,23^, and that aldosterone induces hypertension and vascular dysfunction via CCR5-dependent mechanisms^12^. In addition, we demonstrated that CCL5 stimulates NADPH oxidase 1 (NOX1) activation, positioning CCR5 upstream of redox-dependent vascular injury^12,20^. The present study expands this redox paradigm by identifying mitochondrial dysfunction as a key downstream consequence of CCR5 activation, suggesting that NOX1-derived cytosolic ROS may initiate mitochondrial impairment through redox crosstalk.

A central mechanistic insight of this study is the dissociation between mitochondrial ROS–dependent endothelial dysfunction and ROS-independent vascular hypercontractility. MnTMPyP restored endothelium-dependent relaxation in CCL5-treated vessels, consistent with literature demonstrating that mitoROS reduces nitric oxide bioavailability and contributes to endothelial impairment^25–28^. However, MnTMPyP did not normalize enhanced contractility and increased vasoconstriction in control vessels, suggesting that physiological mitochondrial redox tone contributes to basal modulation of vascular reactivity. In CCL5-treated vessels, hypercontractility appears driven by sustained bioenergetic disruption rather than acute ROS excess. Reduced maximal respiratory capacity and insensitivity to acute mitochondrial uncoupling indicate diminished energetic reserve and persistent membrane potential impairment.

Mitochondrial bioenergetic disruption may influence VSMC contractility through several mechanisms, including altered mitochondrial Ca²⁺ buffering^29^ and enhanced cytosolic Ca²⁺ availability^30,31^, activation of RhoA/Rho kinase–mediated Ca²⁺ sensitization^32,33^, and reduced ATP-sensitive potassium (K_ATP_) channel activity secondary to impaired ATP production^34–36^. These pathways provide mechanistic explanations for persistent contractile enhancement despite ROS scavenging and highlight bioenergetic dysfunction as a distinct therapeutic target.

Although CCR5 deficiency prevented Ang II–induced aortic remodeling, infusion of recombinant CCL5 for 14 days did not elicit structural remodeling in either conduit or resistance vessels. This dissociation suggests that CCR5 signaling is necessary but not sufficient to drive structural remodeling under the present experimental conditions. Ang II activates multiple profibrotic and growth pathways—including mechanical stress– dependent signaling, transforming growth factor-β activation, and broader cytokine networks^37,38^—that are not fully recapitulated by isolated CCL5 exposure. In addition, CCR5 is activated by multiple ligands beyond CCL5, and genetic deletion of CCR5 may blunt a broader chemokine network engaged during Ang II infusion. Longer exposure, higher chemokine burden, or combinatorial stimulation with Ang II–dependent pathways may be required to induce overt structural remodeling.

From a translational perspective, these findings are particularly relevant. CCR5 antagonists such as maraviroc — the first approved small-molecule CCR5 antagonist — are clinically licensed for treatment of CCR5-tropic HIV-1 infection and have well-characterized safety and tolerability profiles based on multiple clinical trials and long-term follow-up studies (e.g., clinical evaluations of maraviroc safety up to 5 years in treatment-experienced patients^39^, 96-week combined safety/efficacy analysis^40^, clinical profile and approval background^41^)^5,39–41^. Our demonstration that CCR5 antagonism prevents CCL5-induced mitochondrial remodeling and mitoROS generation suggests that pharmacologic CCR5 inhibition may not only attenuate inflammatory signaling but also preserve mitochondrial function in the vasculature. Given that mitochondrial dysfunction and oxidative stress are hallmarks of vascular disease in hypertension^9,12^, metabolic syndrome^42^, and atherosclerosis^43^, targeting the CCL5/CCR5 axis could represent a strategy to interrupt a feed-forward inflammatory–mitochondrial injury loop.

Importantly, current antihypertensive therapies primarily target hemodynamic pathways, including the renin–angiotensin system and sympathetic activation. The present findings suggest that adjunctive targeting of chemokine signaling, and mitochondrial bioenergetics may provide additional vascular protection beyond blood pressure reduction alone. In particular, patients with inflammatory or immune-activated hypertensive phenotypes may benefit from modulation of CCR5 signaling. However, because CCR5 plays a central role in immune cell trafficking and host defense^44,45^, therapeutic strategies aimed at chemokine pathway inhibition would require careful patient selection and monitoring to avoid unintended immunological consequences. The identification of a CCR5-dependent mitochondrial mechanism provides a framework for exploring precision-based approaches that target inflammatory–mitochondrial crosstalk while preserving essential immune function.

It is important to highlight some limitations of our work. Although our findings support a NOX1–mitochondrial axis downstream of CCR5, direct molecular dissection of this redox crosstalk requires further investigation. In addition, while short-term CCL5 infusion induced functional impairment without overt structural remodeling, longer-term exposure may produce progressive vascular structural changes.

In conclusion, Ang II amplifies CCL5/CCR5 signaling in the vasculature, and CCR5 activation drives mitochondrial dysfunction characterized by impaired respiratory reserve, disrupted membrane potential, and increased mitochondrial oxidative stress. These mitochondrial alterations underlie endothelial dysfunction and altered vascular contractility. Targeting the CCL5/CCR5 axis and its mitochondrial consequences may represent a translational strategy to mitigate vascular injury in hypertension.

## Acknowledgment

This work was supported by the São Paulo Research Foundation (FAPESP), grant number 2024/14927-3 and NIH-R01HL169202 and startup funds from University of South Alabama to TBN.

## REFERENCES

1. Whelton PK, Carey RM, Aronow WS, et al. 2017 ACC/AHA/AAPA/ABC/ACPM/AGS/APhA/ASH/ASPC/NMA/PCNA Guideline for the Prevention, Detection, Evaluation, and Management of High Blood Pressure in Adults: Executive Summary: A Report of the American College of Cardiology/American Heart Association Task Force on Clinical Practice Guidelines. Circulation. 2018;138(17):e426–e483.

2. Zhang J, Yin Z, Xu Y, et al. Resolvin E1/ChemR23 Protects Against Hypertension and Vascular Remodeling in Angiotensin II-Induced Hypertensive Mice. Hypertension. 2023;80(12):2650–2664.

3. Bader M, Steckelings UM, Alenina N, Santos RAS, Ferrario CM. Alternative Renin-Angiotensin System. Hypertension. 2024;81(5):964–976.

4. Mehta PK, Griendling KK. Angiotensin II cell signaling: physiological and pathological effects in the cardiovascular system. Am J Physiol Cell Physiol. 2007;292(1):C82–97.

5. Jones KL, Maguire JJ, Davenport AP. Chemokine receptor CCR5: from AIDS to atherosclerosis. Br J Pharmacol. 2011;162(7):1453–1469.

6. Zhang Z, Wang Q, Yao J, et al. Chemokine Receptor 5, a Double-Edged Sword in Metabolic Syndrome and Cardiovascular Disease. Front Pharmacol. 2020;11:146.

7. Lv S, Zhou M, Li T, et al. CCL5 promotes angiotensin II-induced cardiac remodeling through regulation of platelet-driven M2 macrophage polarization. Theranostics. 2026;16(2):689–711.

8. Mateo T, Abu Nabah YN, Abu Taha M, et al. Angiotensin II-induced mononuclear leukocyte interactions with arteriolar and venular endothelium are mediated by the release of different CC chemokines. J Immunol. 2006;176(9):5577–5586.

9. Mikolajczyk TP, Nosalski R, Szczepaniak P, et al. Role of chemokine RANTES in the regulation of perivascular inflammation, T-cell accumulation, and vascular dysfunction in hypertension. FASEB J. 2016;30(5):1987–1999.

10. Oliveira de Moraes LH, Beling T, Felix Pimenta G, Bruder-Nascimento T. Chemokine receptors in vascular biology: a review of current evidence, implications, and therapeutic targets for hypertension. Clin Sci (Lond). 2025;139(16):880–895.

11. Singh S, Bruder-Nascimento A, Belin de Chantemele EJ, Bruder-Nascimento T. CCR5 antagonist treatment inhibits vascular injury by regulating NADPH oxidase 1. Biochem Pharmacol. 2022;195:114859.

12. Costa RM, Cerqueira DM, Bruder-Nascimento A, et al. Role of the CCL5 and Its Receptor, CCR5, in the Genesis of Aldosterone-Induced Hypertension, Vascular Dysfunction, and End-Organ Damage. Hypertension. 2024;81(4):776–786.

13. Zorov DB, Juhaszova M, Sollott SJ. Mitochondrial reactive oxygen species (ROS) and ROS-induced ROS release. Physiol Rev. 2014;94(3):909–950.

14. Awata WMC, Alves JV, Costa RM, et al. Vascular injury associated with ethanol intake is driven by AT1 receptor and mitochondrial dysfunction. Biomed Pharmacother. 2023;169:115845.

15. Camacho-Encina M, Booth LK, Redgrave RE, Folaranmi O, Spyridopoulos I, Richardson GD. Cellular Senescence, Mitochondrial Dysfunction, and Their Link to Cardiovascular Disease. Cells. 2024;13(4).

16. Dikalov SI, Nazarewicz RR, Bikineyeva A, et al. Nox2-induced production of mitochondrial superoxide in angiotensin II-mediated endothelial oxidative stress and hypertension. Antioxid Redox Signal. 2014;20(2):281–294.

17. Inagaki S, Suzuki Y, Kawasaki K, Kondo R, Imaizumi Y, Yamamura H. Mitofusin 1 and 2 differentially regulate mitochondrial function underlying Ca(2+) signaling and proliferation in rat aortic smooth muscle cells. Biochem Biophys Res Commun. 2023;645:137–146.

18. Murphy MP. How mitochondria produce reactive oxygen species. Biochem J. 2009;417(1):1–13.

19. Singh S, Bruder A, Costa RM, et al. Vascular Contractility Relies on Integrity of Progranulin Pathway: Insights Into Mitochondrial Function. J Am Heart Assoc. 2025;14(3):e037640.

20. Bruder-Nascimento A, Awata WMC, Alves JV, Singh S, Costa RM, Bruder-Nascimento T. Progranulin Maintains Blood Pressure and Vascular Tone Dependent on EphrinA2 and Sortilin1 Receptors and Endothelial Nitric Oxide Synthase Activation. J Am Heart Assoc. 2023;12(16):e030353.

21. Makrecka-Kuka M, Krumschnabel G, Gnaiger E. High-Resolution Respirometry for Simultaneous Measurement of Oxygen and Hydrogen Peroxide Fluxes in Permeabilized Cells, Tissue Homogenate and Isolated Mitochondria. Biomolecules. 2015;5(3):1319–1338.

22. Lopes RA, Neves KB, Pestana CR, et al. Testosterone induces apoptosis in vascular smooth muscle cells via extrinsic apoptotic pathway with mitochondria-generated reactive oxygen species involvement. Am J Physiol Heart Circ Physiol. 2014;306(11):H1485–1494.

23. Costa RM, Cerqueira DM, Bruder-Nascimento A, et al. Role Of The C-C Motif Chemokine Ligand 5 (CCL5) And Its Receptor, C-C Motif Chemokine Receptor 5 (CCR5) In The Genesis Of Aldosterone-induced Hypertension, Vascular Dysfunction, And End-organ Damage. bioRxiv. 2023.

24. Bruder-Nascimento T, Kress TC, Kennard S, Belin de Chantemele EJ. HIV Protease Inhibitor Ritonavir Impairs Endothelial Function Via Reduction in Adipose Mass and Endothelial Leptin Receptor-Dependent Increases in NADPH Oxidase 1 (Nox1), C-C Chemokine Receptor Type 5 (CCR5), and Inflammation. J Am Heart Assoc. 2020;9(19):e018074.

25. Shah AM, Channon KM. Free radicals and redox signalling in cardiovascular disease. Heart. 2004;90(5):486–487.

26. Jarasuniene D, Simaitis A. [Oxidative stress and endothelial dysfunction]. Medicina (Kaunas). 2003;39(12):1151–1157.

27. Salminen A, Ojala J, Kaarniranta K, Kauppinen A. Mitochondrial dysfunction and oxidative stress activate inflammasomes: impact on the aging process and age-related diseases. Cell Mol Life Sci. 2012;69(18):2999–3013.

28. Ungvari Z, Labinskyy N, Gupte S, Chander PN, Edwards JG, Csiszar A. Dysregulation of mitochondrial biogenesis in vascular endothelial and smooth muscle cells of aged rats. Am J Physiol Heart Circ Physiol. 2008;294(5):H2121–2128.

29. Johnson MT, Benson JC, Pathak T, et al. The airway smooth muscle sodium/calcium exchanger NCLX is critical for airway remodeling and hyperresponsiveness in asthma. J Biol Chem. 2022;298(8):102259.

30. Hajnoczky G, Csordas G, Das S, et al. Mitochondrial calcium signalling and cell death: approaches for assessing the role of mitochondrial Ca2+ uptake in apoptosis. Cell Calcium. 2006;40(5-6):553–560.

31. Xia Y, Li B, Zhang F, et al. Hydroxyapatite nanoparticles promote mitochondrial-based pyroptosis via activating calcium homeostasis and redox imbalance in vascular smooth muscle cells. Nanotechnology. 2022;33(27).

32. Uematsu N, Ogawa K, Tokinaga Y, Tange K, Hatano Y. Sevoflurane inhibits angiotensin II-induced Rho kinase-mediated contraction of vascular smooth muscle from spontaneously hypertensive rat. J Anesth. 2011;25(3):398–404.

33. Hisaoka T, Yano M, Ohkusa T, et al. Enhancement of Rho/Rho-kinase system in regulation of vascular smooth muscle contraction in tachycardia-induced heart failure. Cardiovasc Res. 2001;49(2):319–329.

34. Brayden JE. Functional roles of KATP channels in vascular smooth muscle. Clin Exp Pharmacol Physiol. 2002;29(4):312–316.

35. Tinker A, Aziz Q, Thomas A. The role of ATP-sensitive potassium channels in cellular function and protection in the cardiovascular system. Br J Pharmacol. 2014;171(1):12–23.

36. Jackson WF. Potassium Channels in Regulation of Vascular Smooth Muscle Contraction and Growth. Adv Pharmacol. 2017;78:89–144.

37. Cau SB, Bruder-Nascimento A, Silva MB, et al. Angiotensin-II activates vascular inflammasome and induces vascular damage. Vascul Pharmacol. 2021;139:106881.

38. Rodriguez-Vita J, Sanchez-Lopez E, Esteban V, Ruperez M, Egido J, Ruiz-Ortega M. Angiotensin II activates the Smad pathway in vascular smooth muscle cells by a transforming growth factor-beta-independent mechanism. Circulation. 2005;111(19):2509–2517.

39. Gulick RM, Fatkenheuer G, Burnside R, et al. Five-year safety evaluation of maraviroc in HIV-1-infected treatment-experienced patients. J Acquir Immune Defic Syndr. 2014;65(1):78–81.

40. Hardy WD, Gulick RM, Mayer H, et al. Two-year safety and virologic efficacy of maraviroc in treatment-experienced patients with CCR5-tropic HIV-1 infection: 96-week combined analysis of MOTIVATE 1 and 2. J Acquir Immune Defic Syndr. 2010;55(5):558–564.

41. Lorenzen T. Profile of maraviroc: a CCR5 antagonist in the management of treatment-experienced HIV patients. HIV AIDS (Auckl). 2010;2:151–156.

42. Prasun P. Mitochondrial dysfunction in metabolic syndrome. Biochim Biophys Acta Mol Basis Dis. 2020;1866(10):165838.

43. Cipriani S, Francisci D, Mencarelli A, et al. Efficacy of the CCR5 antagonist maraviroc in reducing early, ritonavir-induced atherogenesis and advanced plaque progression in mice. Circulation. 2013;127(21):2114–2124.

44. Khan IA, Thomas SY, Moretto MM, et al. CCR5 is essential for NK cell trafficking and host survival following Toxoplasma gondii infection. PLoS Pathog. 2006;2(6):e49.

45. Nguyen NZN, Tran VG, Lee S, et al. CCR5-mediated Recruitment of NK Cells to the Kidney Is a Critical Step for Host Defense to Systemic Candida albicans Infection. Immune Netw. 2020;20(6):e49.

