## Supplemental info for "CCR5/CCL5-Dependent Mitochondrial Dysfunction Contributes to Angiotensin II– Induced Vascular Impairment in Mice"

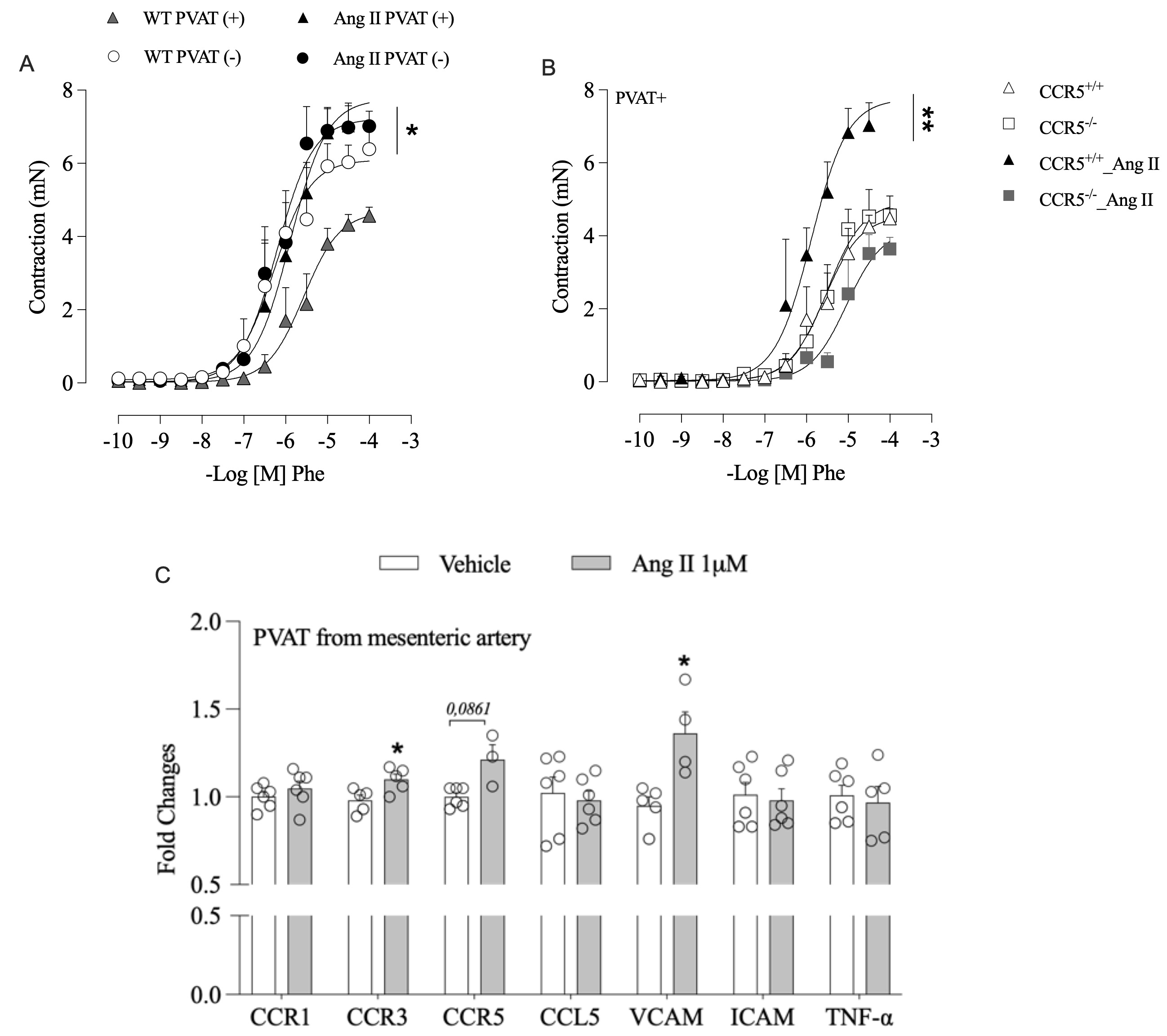
**Supplementary Figure 1. CCR5 contributes to Ang II–induced PVAT dysfunction. Concentration–response curves to phenylephrine (PE) in mesenteric arteries from wild-type (WT) mice with intact perivascular adipose tissue (PVAT+) or with PVAT removed (PVAT−), treated with vehicle (saline) or angiotensin II (Ang II; 490 ng/kg/min for 14 days via osmotic minipump) (A). Concentration–response curves to PE in mesenteric arteries with intact PVAT (PVAT+) from CCR5⁺/⁺ and CCR5⁻/⁻ mice treated with vehicle or Ang II (490 ng/kg/min for 14 days via osmotic minipump) (B). Gene expression of chemokine receptors and inflammatory markers in PVAT isolated from mesenteric arteries of vehicle- or Ang II–treated mice, determined by real-time RT-PCR (C). Data are expressed as mean ± SEM (n = 4–5 per group). Emax and pD₂ values were calculated by nonlinear regression. Concentration–response curves were analyzed using two-way ANOVA followed by Tukey’s post hoc test. Gene expression data were analyzed using Student’s t test or two-way ANOVA, as appropriate.** *P < 0.05 vs. respective vehicle or WT PVAT (+); **P < 0.05 vs. CCR5⁺/⁺, CCR5⁻/⁻, and CCR5⁻/⁻_Ang II.

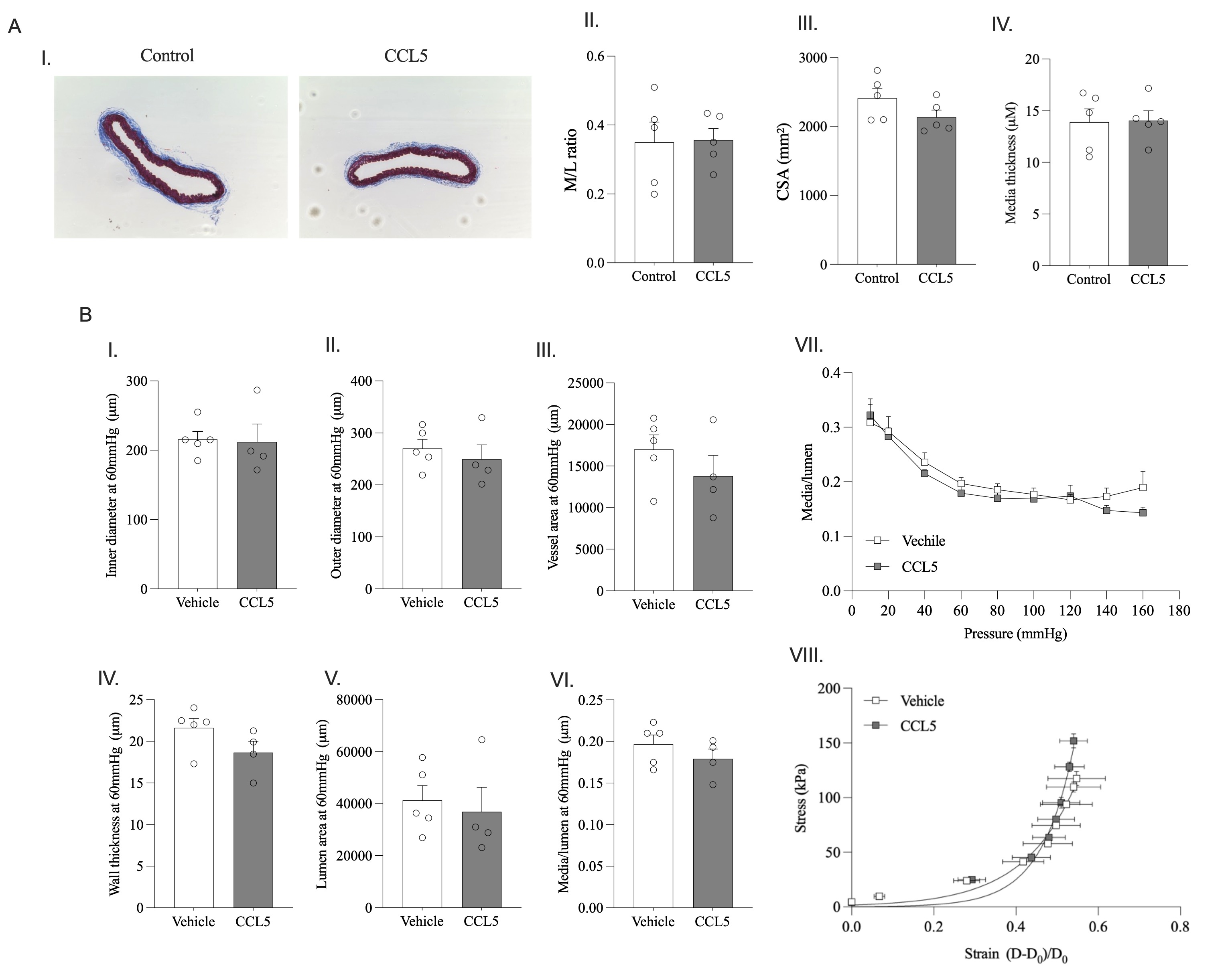

**Supplementary Figure 2. CCL5 does not induce structural or mechanical remodeling in conduit or resistance vessels. Vascular structural and mechanical properties in thoracic aortae (A) and mesenteric arteries (B) from wild-type mice infused with vehicle (saline) or recombinant CCL5 (0.42 ng/day for 14 days via osmotic minipump). (A) Representative Masson’s trichrome–stained sections of thoracic aorta (I). Quantification of media-to-lumen ratio (II), cross-sectional area (CSA) (III), and medial thickness (IV). (B) Structural and biomechanical parameters of mesenteric arteries assessed by pressure myography, including internal diameter (I), external diameter (II), vessel wall area (III), wall thickness (IV), lumen area (V), and media-to-lumen ratio (VI) at 60 mmHg. Media-to-lumen ratio across a range of intraluminal pressures (10–160 mmHg) (VII) and stress–strain relationship (VIII). Data are expressed as mean ± SEM (n = 4–5 per group). Statistical analysis was performed using Student’s t test. No significant differences were observed between vehicle- and CCL5-treated groups.**

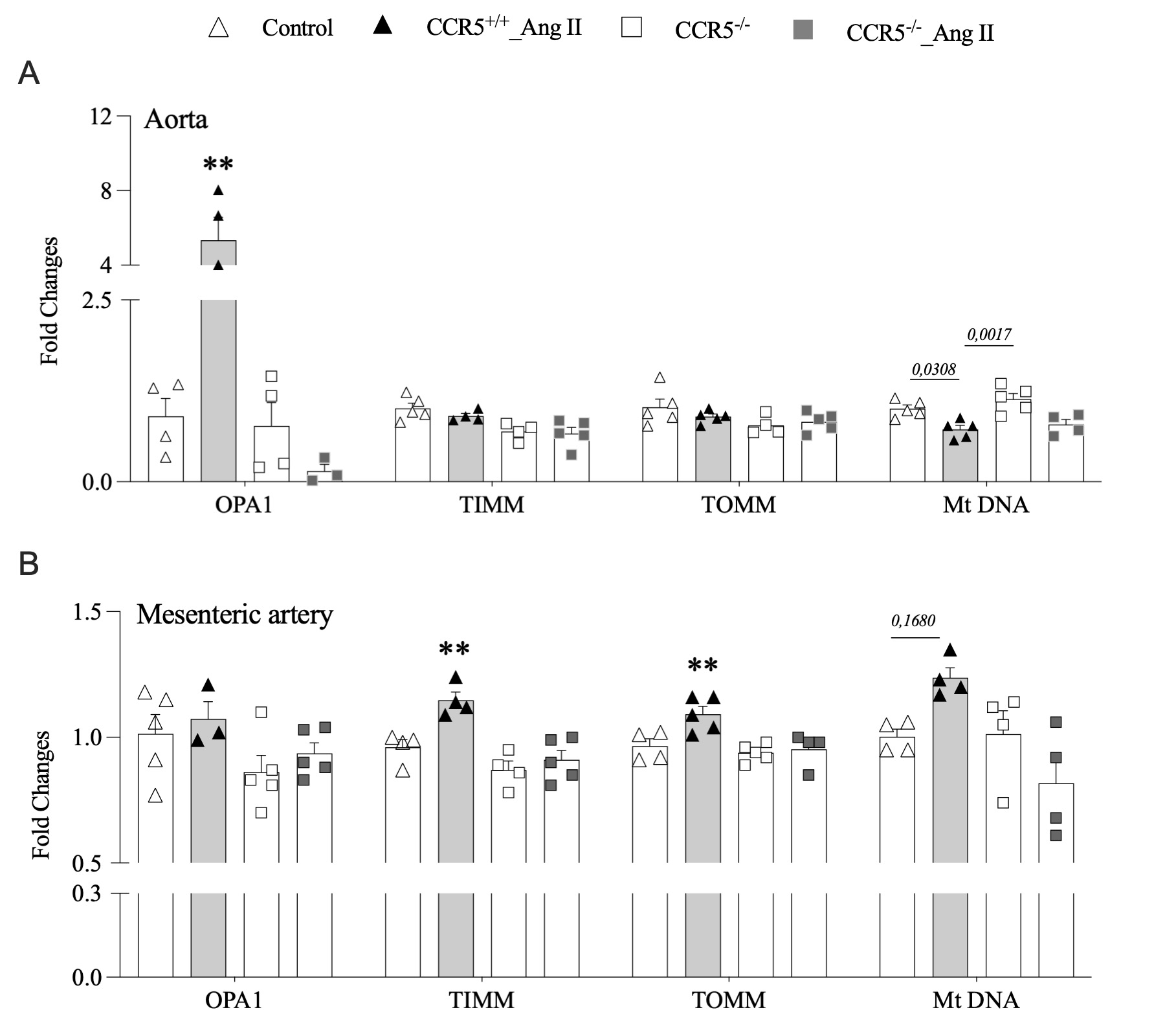

**Supplementary Figure 3. CCR5 deficiency prevents Ang II–induced alterations in mitochondrial-related gene expression.** Gene expression of mitochondrial dynamics and redox-related genes in thoracic aortae (A) and mesenteric arteries (B) from CCR5⁺/⁺ and CCR5⁻/⁻ mice treated with vehicle or angiotensin II (Ang II; 490 ng/kg/min for 14 days via osmotic minipump), determined by real-time RT-PCR. Data are presented as mean ± SEM (n= 3-5). Statistical analysis was performed using two-way ANOVA followed by Tukey’s post hoc test. **P < 0.05 vs. control, CCR5⁺/⁺_Ang II, and CCR5⁻/⁻_Ang II.

**TABLES**

**Table S1:** Sequences of forward and reverse primers used for RT-PCR (Mouse).

| **Target genes** | | Sequence |
| --- | --- | --- |
| CCR1 | FW | GCCAAAAGACTGCTGTAAGAGCC |
|  | RV | GCTTTGAAGCCTCCTATGCTGC |
| CCR2 | FW | GCTGTGTTTGCCTCTCTACCAG |
|  | RV | CAAGTAGAGGCAGGATCAGGCT |
| CCR3 | FW | CCACTGTACTCCCTGGTGTTCA |
|  | RV | GGACAGTGAAGAGAAAGAGCAGG |
| CCR4 | FW | GGACTAGGTCTGTGCAAGATCG |
|  | RV | TGCCTTCAAGGAGAATACCGCG |
| CCR5 | FW | GGTTCCTGAAAGCGGCTGTAAATA |
|  | RV | CTGTTGGCAGTCAGGCACATC |
| CCL5 | FW | AGATCTCTGCAGCTGCCCTCA |
|  | RV | GGAGCACTTGCTGCTGGTGTAG |
| CCL2 | FW | GCTACAAGAGGATCACCAGCAG |
|  | RV | GTCTGGACCCATTCCTTCTTGG |
| ICAM | FW | ATCACATGGGTCGAGGGTTT |
|  | RV | AACCACTGCCAGTCCACATA |
| VCAM | FW | TGACAAGTCCCCATCGTTGA |
|  | RV | ACCTCGCGACGGCATAATT |
| F480 | FW | CGTGTTGTTGGTGGCACTGTGA |
|  | RV | CCACATCAGTGTTCCAGGAGAC |
| IL-6 | FW | CACCCAGAACTTCCATCCACA |
|  | RV | AGCTATGCTCCTCCGTGGCTG |
| IL-16 | FW | CACGCAGACTTCATCCTCCACA |
|  | RV | AGCTATAGTCCATCCGTGCCTG |
| TNF-a | FW | AATGGCCTCCCTCTCATCAG |
|  | RV | CCTAACTGCCCTTCCTCCAT |
| IL1-b | FW | TGACGGACCCCAAAAGATGA |
|  | RV | GCTCTTGTTGATGTGCTGCT |
| iNOS | FW | GAGACAGGGAAGTCTGAAGCAC |
|  | RV | CCAGCAGTAGTTGCTCCTCTTC |
| SOD2 | FW | TAACGCGCAGATCATGCAGCTG |
|  | RV | AGGCTGAAGAGCGACCTGAGTT |
| OPA1 | FW | TCTCAGCCTTGCTGTGTCAGAC |
|  | RV | TTCCGTCTCTAGGTTAAAGCGCG |
| TIMM | FW | AGCTCGAGGCTAGACCATCA |
|  | RV | CAAGCTCTTCCCTTGTCCAG |
| TOMM | FW | AGGAGGGCACYGTCATGTCT |
|  | RV | CTCAAACTCCACACCCACCT |
| MtDNA | FW | CCCCAGCCATAACACAGTATCAAAC |
|  | RV | GCCCAAAGAATCAGAACAGATGC |
| GAPDH | FW | GAGAGGCCCTATCCCAACTC |
|  | RV | TCAAGAGAGTAGGGAGGGCT |

**Table S2**: Sequences of forward and reverse primers used for RT-PCR (Rat).

| **Target genes** | | Sequence |
| --- | --- | --- |
| CCR1 | FW | ACCCCAGGAATTGACCACTG |
|  | RV | AGGCTTTGTTTCTGGGCCTT |
| CCR2 | FW | GCAAAGACCAGAAAAGGGCA |
|  | RV | GGCAGGATCCAAGCTCCAAT |
| CCR3 | FW | ACCTGAGAAGCTAGCCTGTTT |
|  | RV | ACCATCATGTTGCCCAGGAG |
| CCR5 | FW | TGCTCCTGCCATTTGGGATT |
|  | RV | AATGGAAGGCAGCTCTGGTC |
| CCL5 | FW | CTGCTGCTTTGCCTACCTCT |
|  | RV | ATCCCCAGCTGGTTAGGACT |
| IL-6 | FW | CTGCTCTGGTCTTCTGGAGT |
|  | RV | AGAGCATTGGAAGTTGGGGT |
| IL-1b | FW | TTGAGTCTGCACAGTTCCCC |
|  | RV | TCCTGGGGAAGGCATTAGGA |
| iNOS | FW | ATTCAGATCCCGAAACGC |
|  | RV | CCAGAACCTCCAGGCACA |
| COX1 | FW | AGCACATTCGGTGGTGATGT |
|  | RV | GGGTAATCTGGCACACGGAA |
| COX2 | FW | GTGGAAAAGCCTCGTCCAGA |
|  | RV | TCCTCCGAAGGTGCTAGGTT |
| PGC1-a | FW | ATGAGAAGCGGGAGTCTGAA |
|  | RV | GCGGTCTCTCAGTTCTGTCC |
| GAPDH | FW | GAGAGGCCCTATCCCAACTC |
|  | RV | TCAAGAGAGTAGGGAGGGCT |
| FW: Forward; RV: Reverse | | |

**Table S3**: List of antibodies-western blot.

| **Antibody** | **Company** | **Catalog Number** | **Dilution** |
| --- | --- | --- | --- |
| COX-IV | Cell signaling | #4850 | 1:1000 |
| OPA-1 | Cell signaling | #80471 | 1:1000 |
| GAPDH | Cell signaling | #8884 | 1:1000 |
| β-ACTIN | Sigma | #A3854 | 1:20000 |
